# Two tandem mechanisms control bimodal expression of the flagellar genes in *Salmonella enterica*

**DOI:** 10.1101/2019.12.18.881938

**Authors:** Xiaoyi Wang, Santosh Koirala, Phillip D. Aldridge, Christopher V. Rao

**Affiliations:** Department of Chemical and Biomolecular Engineering, University of Illinois at Urbana-Champaign, Urbana, Illinois, United States, 61801; Carl R. Woese Institute for Genomics Biology, University of Illinois at Urbana-Champaign, Urbana, Illinois, United States, 61801; Centre for Bacterial Cell Biology, Newcastle University, Newcastle upon Tyne, United Kingdom

## Abstract

Flagellar gene expression is bimodal in *Salmonella enterica*. Under certain growth conditions, some cells express the flagellar genes whereas others do not. This results in mixed populations of motile and non-motile cells. In the present study, we found that two independent mechanisms control bimodal expression of the flagellar genes. One was previously found to result from a double negative-feedback loop involving the flagellar regulators YdiV and FliZ. This feedback loop governs bimodal expression of class 2 genes. In this work, a second mechanism was found to govern bimodal expression of class 3 genes. In particular, class 3 gene expression is still bimodal even when class 2 gene expression is not. Using a combination of experimental and modeling approaches, we found that class 3 bimodalilty results from the σ^28^-FlgM developmental checkpoint.

**IMPORTANCE:** Many bacterial use flagella to swim in liquids and swarm over surface. In *Salmonella enterica*, over fifty genes are required to assemble flagella. The expression of these genes is tightly regulated. Previous studies have found that flagella gene expression is bimodal in *S. enterica*, which means that only a fraction of cells express flagellar genes and are motile. In the present study, we found that two separate mechanisms induce this bimodal response. One mechanism, which was previously identified, tunes the fraction of motile cells in response to nutrients. The other results from a developmental checkpoint that couples flagellar gene expression to flagellar assembly. Collectively, these results further our understanding of how flagellar gene expression is regulated in *S. enterica*.

## INTRODUCTION

Many bacteria can switch between motile and non-motile states. Food is often a key factor in determining whether these bacteria are motile or not. For example, many bacteria are motile only when grown in nutrient-limited media; others are motile only when grown in nutrient-rich media. *Salmonella enterica* is an example of the latter. This bacterium employs flagella to swim in liquids (1). Previous studies have shown that nutrients induce the expression of the flagellar genes in *S. enterica* (2). The individual bacteria, however, do not all respond the same to nutrients. Rather, nutrients tune the relative fraction of motile and non-motile cells within the population (3). These mixed populations indicate that the response to nutrients is bimodal, where two otherwise identical cells can exhibit an entirely different response to the same nutrient concentrations.

Multiple studies have observed bimodal expression of the flagellar genes in *S. enterica* (3–9). As a brief background, the flagellar promoters can be grouped into three hierarchical classes based on how they are temporally activated (10, 11). A single class 1 promoter controls the expression of the master flagellar regulator, the FlhD_4_C_2_ complex (12). FlhD_4_C_2_ in turn activates class 2 promoters (13). These promoters control the expression of the genes encoding the hook-basal-body (HBB) proteins and two key regulators. One is the alternate sigma factor σ^28^ (FliA), which activates expression of the class 3 promoters. These promoters control expression of the genes encoding the filament, motor, and chemotaxis proteins (14). The other is the anti-sigma factor FlgM (15). Prior to completion of the HBB, FlgM binds σ^28^ and prevents it from activating class 3 promoters. Upon completion of the HBB, FlgM is secreted from the cell, freeing σ^28^ to activate class 3 promoters (16). This mechanism provides a developmental checkpoint, ensuring that class 3 genes are expressed only when functional HBB’s are built. It is also thought to provide a sensing mechanism enabling *S. enterica* to control flagellar abundance (17–21).

A number of additional flagellar proteins are known to regulate flagellar gene expression in *S. enterica* (18, 22–25). In the context of this study, Koirala and coworkers (3) previously demonstrated that a double-negative feedback loop involving two regulatory proteins, FliZ and YdiV, governs bimodal expression of class 2 genes in response to nutrients. YdiV represses class 2 and 3 gene expression by binding to the FlhD subunit of the FlhD_4_C_2_ complex and then promoting its degradation by the protease ClpXP (2, 26). In addition, YdiV prevents the FlhD_4_C_2_ complex from binding to and activating class 2 promoters (2, 26). YdiV also governs the nutrient response: nutrients inhibit expression of YdiV by an unknown mechanism (2). When nutrient concentrations are high, YdiV expression is low, thus freeing FlhD_4_C_2_ to activate class 2 promoters. Conversely, when nutrients concentrations are low, YdiV expression is high, thus preventing FlhD_4_C_2_ from activating class 2 promoters. FliZ, expressed from the hybrid class 2/3 *fliAZ* promoter, activates class 2 and 3 gene expression by inhibiting expression of YdiV, at both the transcriptional and translational level (3, 27). YdiV and FliZ participate in a double-feedback loop, where YdiV indirectly represses expression of FliZ through FlhD_4_C_2_, and FliZ directly represses expression of YdiV (27). As a consequence, two stable expression states are possible: one where YdiV concentrations are high and FliZ concentrations are low; and the other where YdiV concentrations are low and FliZ concentrations are high. In support of this mechanism, only a single expression state (i.e. monostable expression) for class 2 genes is observed when this feedback loop is broken, for example by deleting *fliZ* or *ydiV* (3).

A separate mechanism appears to govern bimodal expression of class 3 genes in *S. enterica*, because class 3 gene expression is still bimodal in a Δ*fliZ* mutant (5, 8). While these previous studies did not investigate the nutrient response per se, they nonetheless demonstrated that the YdiV-FliZ feedback loop does not cause class 3 bimodalilty. In this study, we investigated the mechanism governing the bimodal expression of class 3 genes. In support of previous work, we found that the mechanism is different than the one governing bimodal expression of class 2 genes. Further, we found that it results from the σ^28^-FlgM developmental checkpoint. In the process, our data explain a number of previous results and further our understanding of how flagellar gene expression is regulated in *S. enterica*.

## RESULTS

### FliZ is not does not govern bimodal expression of class 3 genes

We measured the response of class 2 and class 3 promoters to nutrients in single cells using flow cytometry. The goal of these experiments was to determine whether the responses of these two promoter classes were coupled. In particular, would we observe cells where class 2 promoters were active and class 3 promoters inactive? Or, would we only observe cells where both promoter classes were either active or inactive? We eliminated the possibility of observing cells where the class 2 promoters were inactive and the class 3 promoters active from the outset given the known hierarchy among the promoter classes. We further note that previous experiments only measured the response of a single promoter and thus could not be used to examine coupling.

To measure expression from both promoter classes, we created transcriptional fusions of the class 2 *flhB* promoter to the red fluorescent protein mCherry (28) and the class 3 *fliC* promoter to the yellow fluorescent protein Venus (29). These transcriptional reporters were then integrated single copy into the *araB* gene and λ attachment site, respectively. This design enabled us to measure expression from both promoters in single cells using flow cytometry. The cells were grown to late exponential phase in Vogel-Bonner medium E supplemented (30) with 0.2% glucose and various concentrations of yeast extract, where the latter served as the inducing nutrient, prior to analysis by flow cytometry.

The response to yeast extract is shown in Figure 1A. Consistent with previous studies, we observed two co-existing populations at intermediate (0.2-1%) yeast extract concentrations, one where both promoters were inactive and the other where both promoters were active. We further found that the activities of both promoters were coupled: in cells where the class 2 *flhB* promoter was active, the class 3 *fliC* promoter was also active. We did not observe any population where the class 2 *flhB* promoter was active and the class 3 *fliC* promoter was inactive, which would correspond to the upper left quadrant in the panels of Figure 1A. Collectively, these results demonstrate that the responses of these two promoters are tightly coupled. These results are not particularly surprising given the transcriptional hierarchy within the flagellar gene network. One aspect not considered in the present study was the temporal response, where we would expect activation of the class 2 promoters to precede activation of the class 3 promoters during early exponential phase, with an intervening lag in between (17).

**Figure 1.**
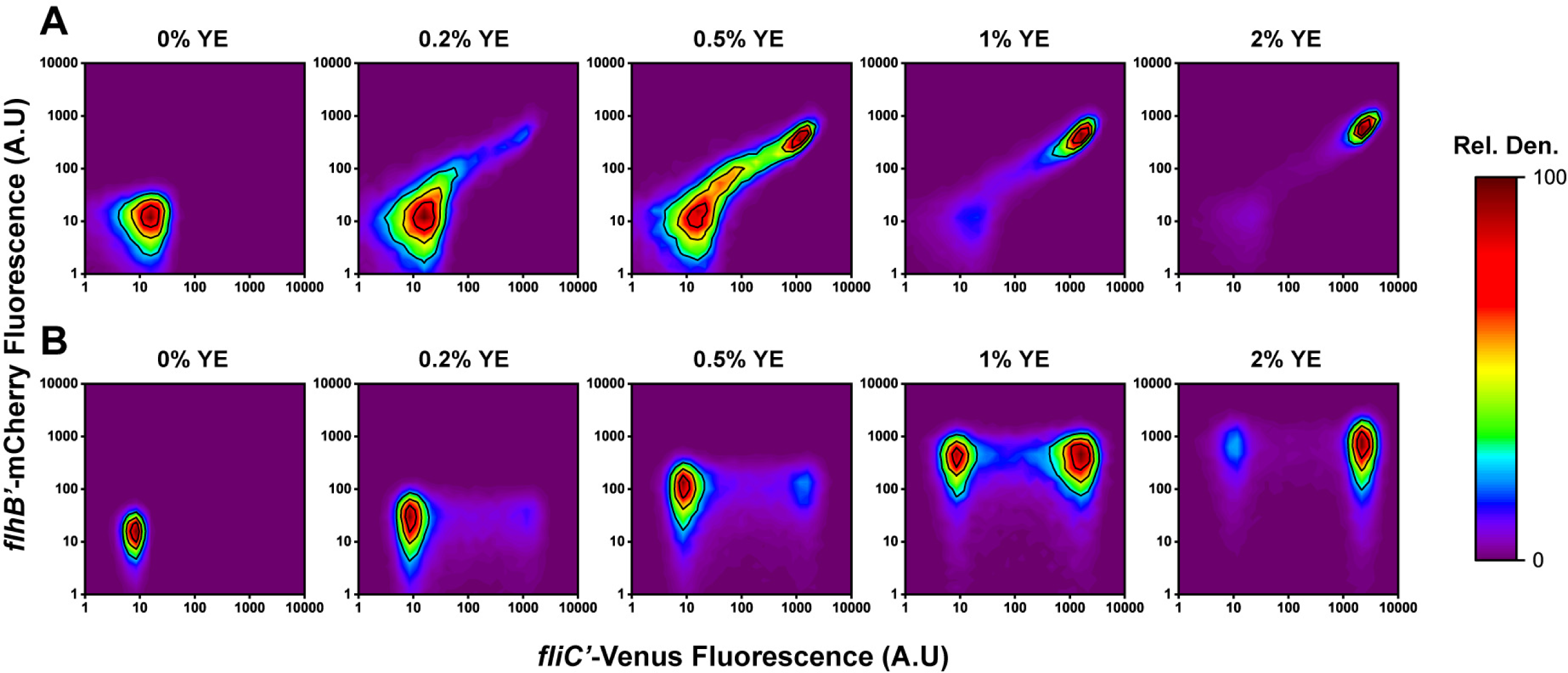
Response of class 2 *flhB* and class 3 *fliC* promoters for wild type (A) and *fliZ* mutant (B) to nutrients (yeast extract) using two-color flow cytometry as determined using single-copy transcriptional fusions of the class 2 *flhB* gene and the class 3 *fliC* gene to the fluorescent proteins mCherry and Venus, respectively. The heatmaps show the relative number (Rel. Den.) of cells exhibiting different levels of *flhB* and *fliC* promoter activity. **Figure S1** provides histograms for the same data.

We next investigated the response of a Δ*fliZ* mutant. Previous studies have shown that the bimodal response of class 2 promoters but not class 3 promoters was eliminated in this mutant (5, 8). These results are confirmed in Figure 1B, where we observed a homogeneous response to nutrients for the class 2 *flhB* promoter and a bimodal response for the class 3 *fliC* promoter. These results clearly demonstrate that separate mechanisms govern the bimodal response of class 2 and class 3 promoters, because we can eliminate it for one promoter class but not for the other.

### Class 3 gene expression is monostable in Δ*ydiV* mutant

FliZ participates with YdiV in a double-negative feedback loop. Furthermore, YdiV governs the nutritional response: nutrients inhibit the expression of YdiV, which in turn inhibits the expression of class 2 genes through FlhD_4_C_2_. Previous studies have shown that class 2 promoters are active (ON state) in a Δ*ydiV* mutant irrespective of whether yeast extract is added. As shown in Figure 2, the same behavior was also observed for the class 3 *fliC* promoter, where this promoter was found to be active (ON state) in all cells. These results are again expected, because the activities of these two promoter classes are coupled (Figure 1). We do note that there was a small population of Δ*ydiV* mutants where the promoters exhibited intermediate levels of expression, distinct from those in the ON state. The origin of this behavior is not known, though others have also observed this intermediate activity state (3, 8).

**Figure 2.**
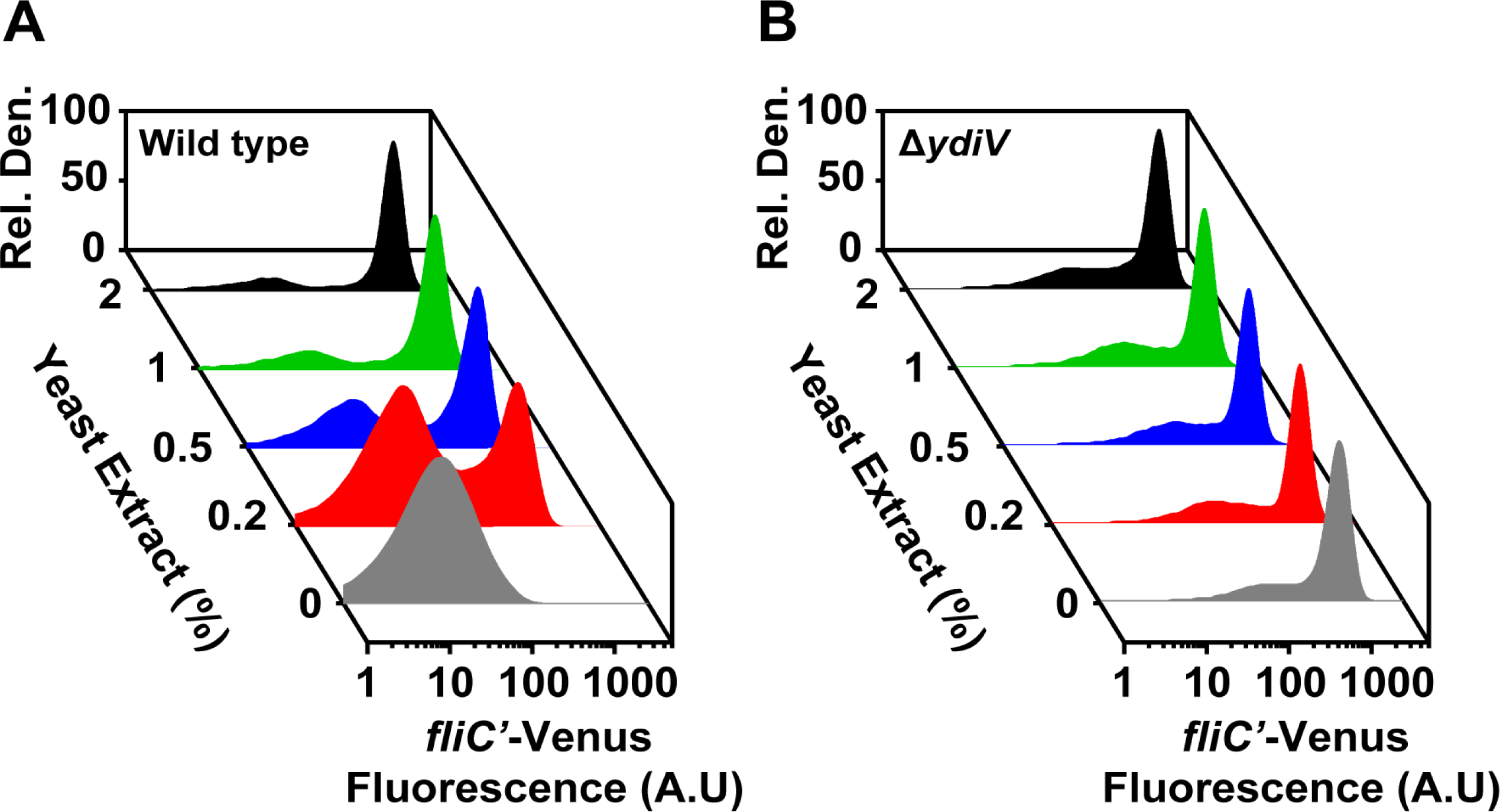
Response of class 3 *fliC* promoters to nutrients (yeast extract) in the wild type (A) and a Δ*ydiV* mutant (B) using flow cytometry as determined using single-copy transcriptional fusion to the fluorescent protein Venus. The histograms show the relative number (Rel. Den.) of cells exhibiting different levels of *fliC* promoter activity.

### FlgM is necessary for class 3 bimodalilty

σ^28^ and FlgM are the principal regulators of class 3 gene expression. We first tested how FlgM affects the expression of class 3 genes by exploring the behavior of a Δ*flgM* mutant. As shown in Figure 3A, the class 3 *fliC* promoter still exhibited a bimodal response to nutrients in a Δ*flgM* mutant. These results do not establish whether FlgM is necessary for class 3 bimodalilty. The reason is that class 2 gene expression is bimodal in a Δ*flgM* mutant (3). As a consequence, class 3 gene expression will also be bimodal in a Δ*flgM* mutant, irrespective of whether there is a separate mechanism for bistablity due to the transcriptional hierarchy among the promoter classes.

**Figure 3.**
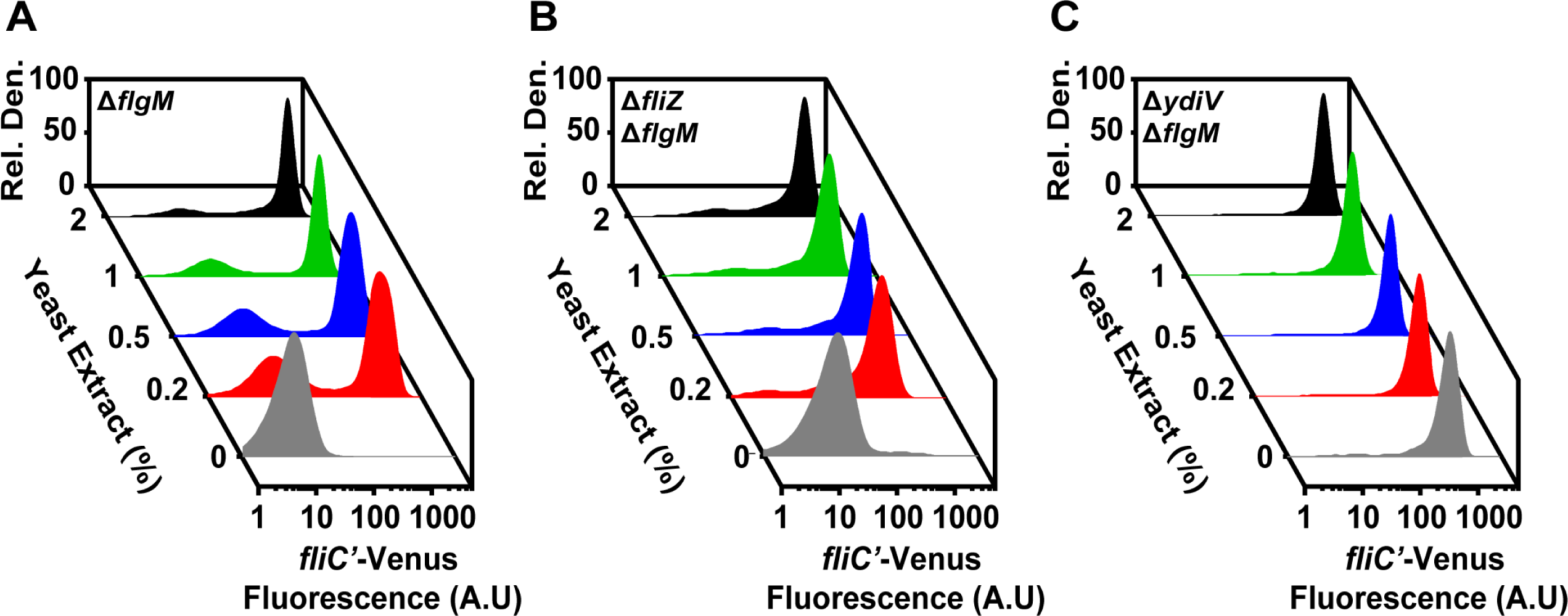
Response of class 3 *fliC* promoter to nutrients (yeast extract) in Δ*flgM* mutant (A), Δ*fliZ* Δ*flgM* mutant (B) and Δ*ydiV* Δ*flgM* (C) mutant using flow cytometry as determined using single-copy transcriptional fusion to the fluorescent protein Venus. The histograms show the relative number (Rel. Den.) of cells exhibiting different levels of *fliC* promoter activity.

To determine whether FlgM is necessary for class 3 bimodalilty, we tested the response of a Δ*fliZ* Δ*flgM* mutant, because class 2 gene expression is unimiodal in this mutant (3). As shown in Figure 3B, the class 3 *fliC* promoters in a Δ*fliZ* Δ*flgM* mutant exhibited a homogenous response to nutrients. We also investigated the response of a Δ*ydiV* Δ*flgM* mutant as a control (Figure 3C). The response in this case was similar to a Δ*ydiV* mutant (Figure 2), where all cells were in the ON state. The only difference is that we observed far less cells in the intermediate expression state. As noted above, we cannot explain this intermediate state. That said, the number of cells in this state was reduced when gene expression was further enhanced due to loss of *flgM*.

### Modeling predicts that the σ^28^-FlgM developmental checkpoint is sufficient to induce class 3 bimodalilty

The results in Figure 3 demonstrate that FlgM is necessary for class 3 bimodalilty in a Δ*fliZ* mutant. These results also suggest that the mechanism most likely involves σ^28^, because FlgM regulates flagellar gene expression by sequestering σ^28^. One hypothesis is that class 3 bimodalilty results from the σ^28^-FlgM developmental checkpoint. To explore this hypothesis, we constructed a simple mathematical model of this checkpoint that relates the concentration of free σ^28^ to the FlgM secretion rate (Figure 4A). This model is a simplified version of a previously published model of flagellar gene regulation (19), in the sense that it focuses only on the σ^28^-FlgM checkpoint (details provided in the methods section).

**Figure 4.**
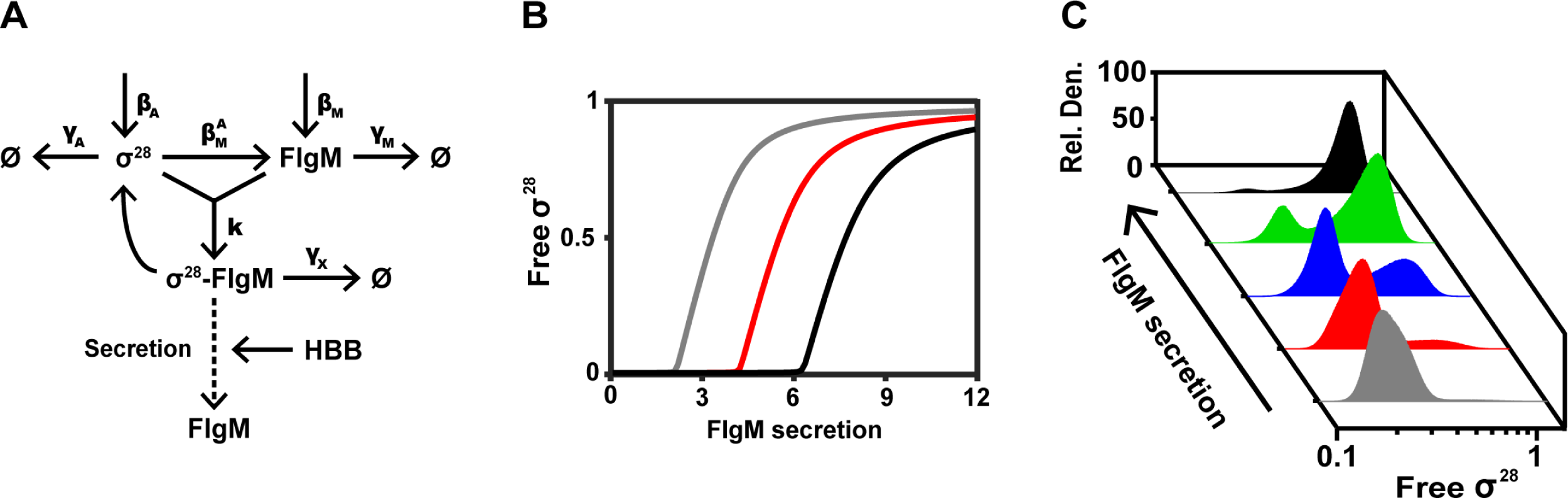
**A.** Key components in the flagellar network that govern class 3 bimodality as described in the mathematical model. **B.** Model predicts that the concentration of free σ^28^ exhibits a sharp threshold with respect to the FlgM secretion rate (parameter *k*_*s*_ in the model). Parameter values: *β*_*A*_ = 1, *γ*_*A*_ = 1, *k* = 10^5^, *K*_*s*_ = 0.5, 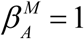, *β*_*M*_ = 1, and *γ*_*X*_ = 0.1. The different curves show how the threshold is determined by the expression of FlgM from the class 2 *flgA* promoter (the parameter *β*_*M*_ in the model: gray curve, *β*_*M*_ = 3; red curve, *β*_*M*_ = 5; black curve, *β*_*M*_ = 7). **C.** Model predicts bimodal distribution of free σ^28^ concentrations within the population. As the FlgM secretion rate increases (parameter *k*_*s*_ in the model), the population shifts from an OFF state to and ON state. The histograms show the relative number (Rel. Den.) of simulated cells with different concentrations of free σ^28^. The parameters are the same as before with *β*_*M*_ = 5. See Materials and Methods for further details.

A representative response is shown in Figure 4B. A critical feature of this response is presence of a threshold. Below this threshold secretion rate, there is no free σ^28^ in the cell – all is bound to FlgM. Only when the FlgM secretion rate exceeds this threshold does the response become hyperbolic. Two mechanisms are responsible for this threshold, which underlies the developmental checkpoint. The first is that σ^28^ induces the expression of *flgM*, which is under the control of both a class 2 and class 3 promoter. This negative feedback loop ensures that sufficient FlgM is produced to effectively sequester any free σ^28^ in the absence of secretion. The second is that the binding of σ^28^ and FlgM is effectively irreversible, with a half-life of approximately one hour (31). This means that if the concentration of FlgM exceeds σ^28^, then all of the σ^28^ will be bound to FlgM. Together these two mechanisms ensure that there is no free σ^28^ in the cell in the absence of secretion. Indeed, this what we observe experimentally (**Figure S2**). However, if the secretion rate is sufficiently high, such that the cell is pumping FlgM out of the cell at a rate faster than it is being produced, then σ^28^ is free to activate the class 3 promoters.

The secretion rate is expected to be proportional to the number of functional HBBs in the cell. As the flow-cytometry data show (Figure 1A), there is significant variability in gene expression among different cells even in the absence of bimodalilty. This means that at intermediate expression states (corresponding to intermediate yeast extract concentrations), some cells may not build enough HBBs to exceed the secretion threshold for inducing class 3 gene expression whereas others will. If the response is sufficiently sharp, then this will suffice in generating class 3 bimodalilty even when distribution of HBBs is homogeneous. To test this hypothesis, we simulated the model assuming that secretion rate was variable within individual cells. We then varied the mean secretion rate, assuming it was homogenously distributed in the population with fixed variance, to mimic the effect of HBB variability. All other model parameters were fixed. As shown in Figure **4C**, variability in the secretion rate is sufficient to generate bimodalilty. Such a mechanism could explain class 3 bimodalilty in a Δ*fliZ* mutant.

To test this prediction, we first replaced the native *fliA* promoter with an anhydrotetracycline-inducible one (P_*fliA*_::*tetRA*). The goal here was to decouple *fliA* expression from the other flagellar genes. When we tested this promoter in a Δ*fliZ*Δ*flgM* mutant, we observed a homogenous response to anhydrotetracycline (aTc) as expected (Figure 5A). In particular, higher σ^28^ expression is expected to result in higher class 3 gene expression. We next explored this promoter in a Δ*fliZ* mutant (Figure 5B). In these experiments, we used yeast extract to tune the expression of the class 2 genes and, indirectly, the rate of FlgM secretion. At low yeast extract concentrations, the class 3 *fliC* promoter was effectively off. This would correspond to the scenario where the secretion rate is below the threshold. However, when the concentrations of yeast extract were increased, we observe the emergence of a second population, corresponding the class 3 ON state. This corresponds to the scenario where some cells exceed the threshold (ON state) and others do not (OFF state).

**Figure 5.**
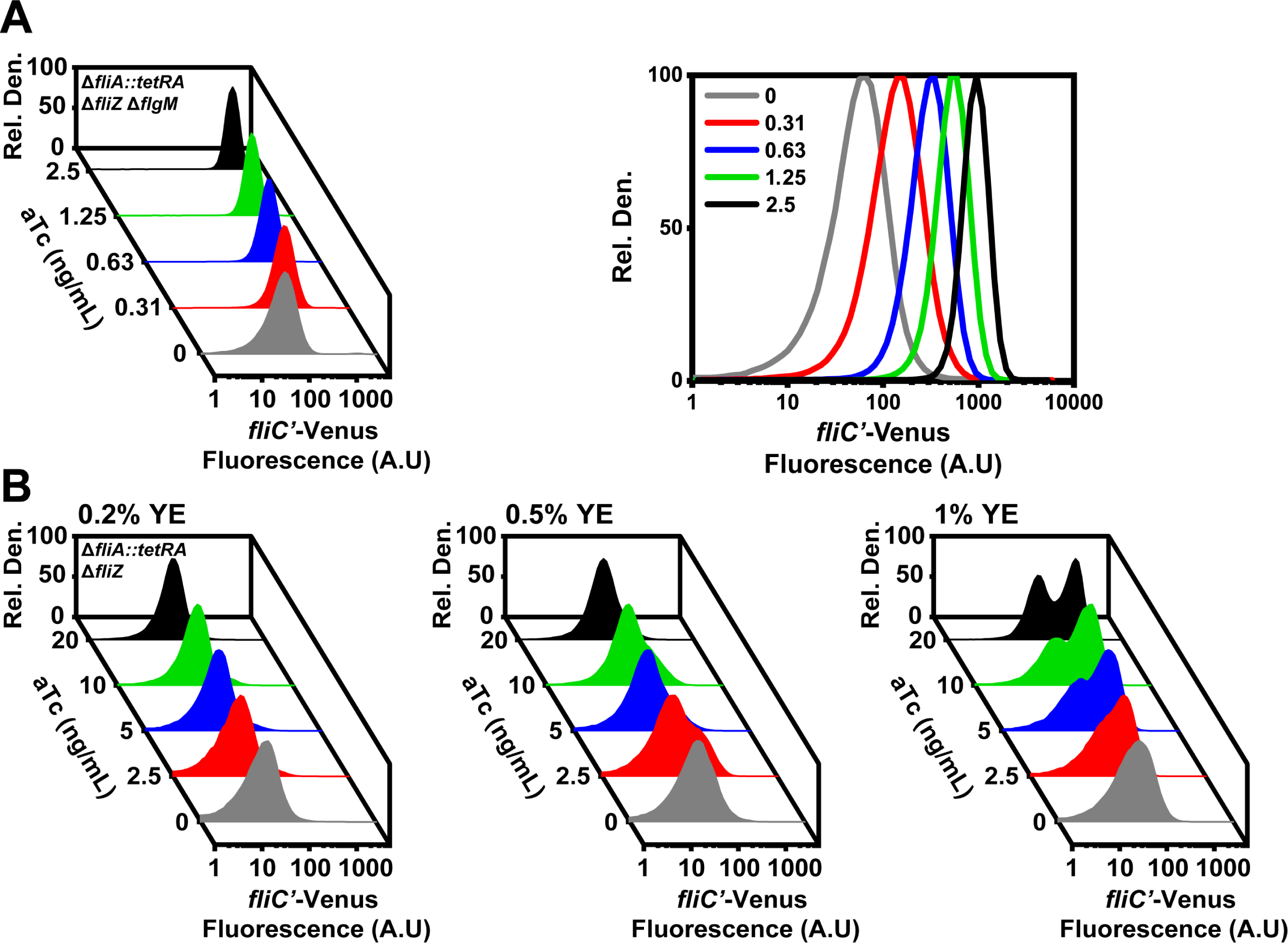
**A.** Response of class 3 *fliC* promoter is monostable with an aTc inducible Δ*fliA*::*tetRA* promoter in a Δ*fliZ* Δ*flgM* mutant. Right panel shows the same data plotted in two dimensions. **B.** Response of class 3 *fliC* promoter with an aTc inducible Δ*fliA*::*tetRA* promoter in a Δ*fliZ* mutant in various yeast extract concentrations. Response was determined using flow cytometry determined using single-copy transcriptional fusion to the fluorescent protein Venus. The histograms show the relative number (Rel. Den.) of cells exhibiting different levels of *fliC* promoter activity.

One limitation of these experiments is that yeast extract represses the tetracycline promoter at high concentrations (>1% yeast extracted) (3), thereby limiting the range of concentrations that can be tested. Therefore, we next replaced native *flgM* promoter with an aTc-inducible one (P_*flgM*_::*tetRA*). In the absence of aTc, we observed two populations, likely due to leaky expression from the *tetRA* promoter (Figure 6). However, when the concentration of aTc was increased, the fraction of cells in the ON state decreased. This was most pronounced at low yeast extract concentrations, where the rate of FlgM secretion is low. At higher yeast extract concentrations, corresponding to higher secretion rates, higher concentrations of aTc were required to reduce the fraction of cells in the ON state.

**Figure 6.**
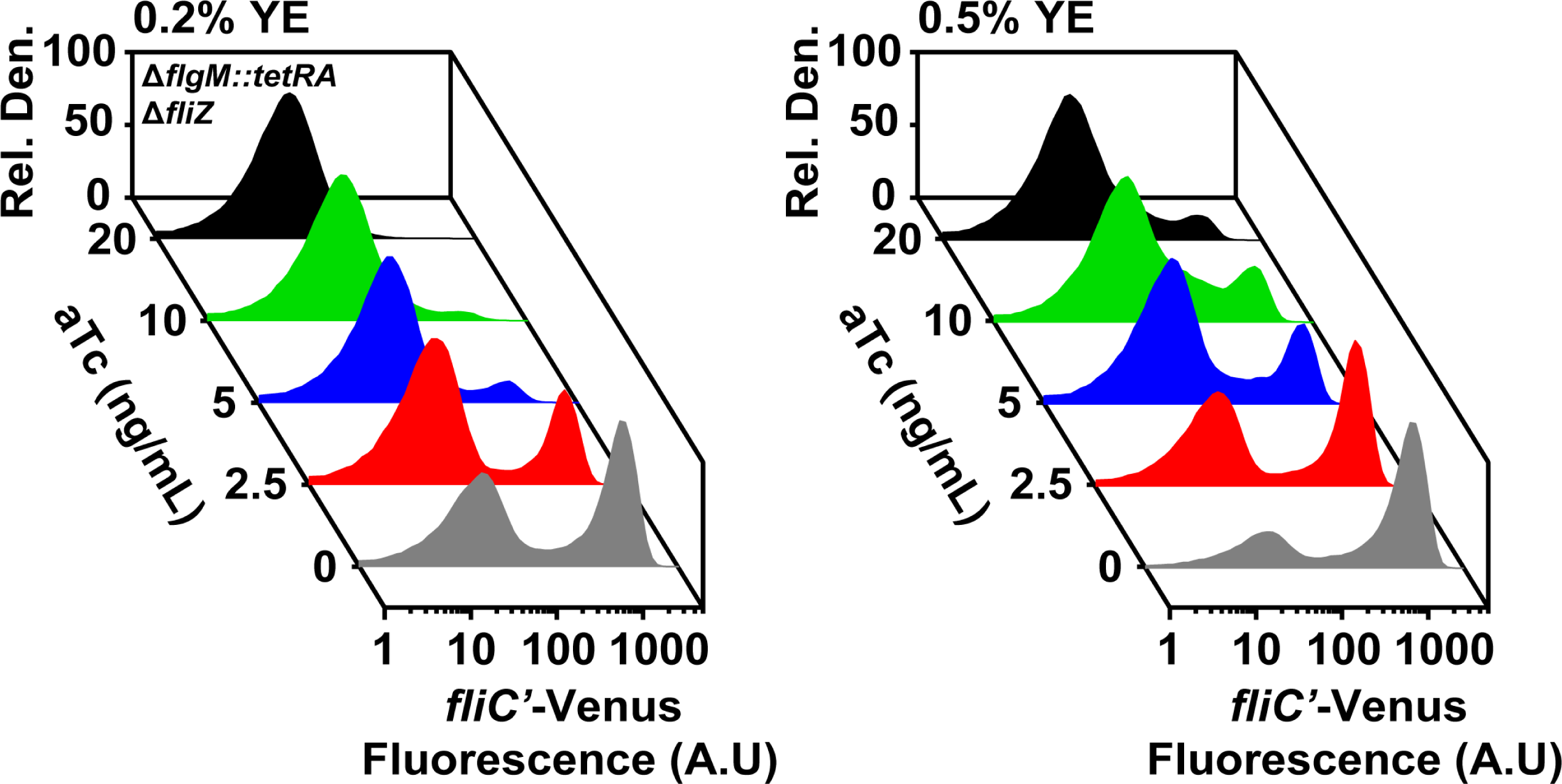
Response of class 3 *fliC* promoter with an aTc inducible Δ*flgM*::*tetRA* promoter in a Δ*fliZ* mutant in 0.2% (left) and 0.5% (right) yeast extract using flow cytometry as determined using single-copy transcriptional fusion to the fluorescent protein Venus. The histograms show the relative number (Rel. Den.) of cells exhibiting different levels of *fliC* promoter activity.

As a further test of our model, we measured class 3 gene expression in a ΔHBB (Δ*flgG-J*) mutant at varying concentrations of yeast exact. This mutant does not build function HBBs and thus is incapable of FlgM secretion. We would expect no class 3 flagellar expression. Consistent with our hypothesis, cells were not able to activate the class 3 gene expression as σ^28^ exists completely in the σ^28^-FlgM complex (**Figure S2**). In the absence of secretion, any σ^28^ produced would be immediately sequestered by FlgM, a key assumption in our model explain the class 3 bimodalilty.

Collectively, these experiments support a number of key model predictions, namely that the class 3 bimodalilty arises from the sharp threshold imposed by the σ^28^-FlgM checkpoint. In other words, our data suggest that the cells need to build a minimum number of HBB’s in order to trigger this checkpoint and activate class 3 gene expression. At intermediate levels of flagellar gene expression, some cells will not have built a sufficient of HBB’s to trigger the checkpoint whereas other will have. Such a mechanism is consistent with our data and would explain class 3 bimodalilty.

## DISCUSSION

Flagellar gene expression is bimodal in *S. enterica* (3–9). Under certain growth conditions, some cells express the flagellar genes whereas others do not. This results in mixed populations of motile and non-motile cells. Nutrients were previously found to specify the fraction of motile cells within the population (3). Whether nutrients alone specify the motile fraction is not presently known. In the present study, we found that two independent mechanisms induce the bimodal expression of the flagellar genes. One induces the bimodal expression of the class 2 genes in response to nutrient concentrations. This was previously found to result from a double-negative feedback loop involving FliZ and YdiV (3). The other induces the bimodal expression of the class 3 genes. The key finding in the present work is that class 3 bimodalilty results from the σ^28^-FlgM checkpoint.

Stewart and coworkers proposed that motility is bimodal in *S. enterica* because it generates mixed populations of invasive and non-invasive cells due to the coupling of motility and virulence i (6, 32, 33). This model, however, does not explain why *S. enterica* employs two mechanisms to induce bimodal expression of the flagellar genes when one alone would suffice. Among the two mechanisms, the FliZ-YdiV feedback loop is clearly dominant, because it specifies the likelihood that an individual cell will be motile or not in response to external nutrient concentrations (and possibly other factors as well). Class 3 bimodalilty, on the other hand, appears to ensure that cells express class 3 genes only when they have built a sufficient number of HBBs. Such a mechanism would efficiently manage resources within the cell, ensuring that cells express class 3 genes only when needed. The effect is masked in the wild type under the conditions explored in this study, because the FliZ-YdiV feedback loop ensures that motile cells builds a sufficient number of HBBs to exceed the threshold. It may, however, play a dynamic role and shut off class 3 gene expression when, for example, a daughter cell inherits too few flagella and only turns it on when sufficient numbers are built. Alternatively, class 3 bimodalilty may simply be a consequence of the sharp threshold imposed by the σ^28^-FlgM checkpoint, one that only manifests itself in mutants with reduced expression of the flagellar genes.

We note that this model extends a previous one proposed for the σ^28^-FlgM checkpoint (19), where it was proposed that it continuously regulates class 3 gene expression in response to HBB abundance using FlgM secretion as proxy signal. That model also predicted that the threshold is not sharp. However, it was based on population-level measurement of gene expression, which lack the resolution necessary to capture phenomena such as bimodalilty. The present study suggests that the threshold is indeed sharp, as this alone explain class 3 bimodalilty (Figure 4). In addition, the original model also predicted that completion of more than one HBB may be necessary to induce class 3 gene expression. The present study supports this claim, because it would explain why some cells exceed the threshold and others do not. If only a single HBB was necessary, then it is unlikely that we clearly observe two populations because the threshold would be more easily exceed (Figure 1).

We also note that the present analysis was limited to steady-state exponential growth. Others have previously explored the temporal dynamics of flagellar gene expression at single-cell resolution (5, 8, 34) and identified a number of transient phenomena that cannot be explained by the working model developed in the present study. Many factors are known to affect the dynamic response of flagellar gene expression. While some have been explored in the past (17–19), these previous studies did not consider heterogeneity among individual cells.

In conclusion, we have demonstrated that the flagellar gene network encodes two mechanisms for bimodal gene expression, one controlling the class 2 genes and the other controlling the class 3 genes. In the process, we have furthered our understanding of how this complex gene network is regulated in *S. enterica*. Our results also emphasize the need to measure flagellar gene expression at single-cell resolution, because bulk assays miss much of the complexity of this regulation. As discussed above, many questions still remain. One concerns the mechanism for nutrient sensing. In particular, this sensing mechanism does not appear to respond to single nutrient but rather the general nutrient/energetic state of the cell (3). It may also respond to other signals as well. Second, additional mechanisms are known to regulate the dynamics of flagellar gene expression. How these regulatory mechanisms manifest themselves at single-cell resolution is still not known. Finally, we still do not know the rates of switching between the non-motile and motile states or whether these transitions are reversible during different phases of growth. More work is needed to answer these questions.

## MATERIALS AND METHODS

### Media and growth conditions

All experiments were performed at 37°C in Vogel-Bonner minimal E (VBE) medium (200 mg/l MgSO_4_.7H_2_O, 2 g/l citric acid monohydrate, 10 g/l anhydrous K_2_HPO_4_ and 3.5 g/l NaNH_4_PO_4_) (30) supplemented with 0.2% glucose and yeast extract at the specified concentrations. Luria-Bertani medium (10 g/l tryptone, 5 g/l yeast extract, 10 g/l NaCl) was used for strain construction. Strains containing the plasmids pKD46, pCP20, and pINT-ts were grown at 30°C. Antibiotics were used at following concentrations: ampicillin at 100 μg/ml, chloramphenicol at 20 μg/ml and kanamycin at 40 μg/ml.

### Bacterial strains and plasmid construction

All strains (Table 1) are derivatives of *S. enterica* serovar Typhimurium 14028 (American Type Culture Collection). The Δ*flhDC* (region 2032540 to 2033471), Δ*flgM* (region 1215209 to 1215502), Δ*fliZ* (region 2055542 to 2056093), Δ*ydiV* (region 1432774 to 1433487), and ΔHBB (Δ*flgG-J*, region 1261788 to 1265393) mutants were constructed using the method of Datsenko and Wanner (Datsenko & Wanner, 2000). The integrated cassettes were then moved to a clean wild-type background by P22 transduction prior to removal of the antibiotic marker with pCP20. Using the same method, the promoters P_*fliA*_ (region 2057139 to 2056887) and P_*flgM*_ (region 1215584 to 1215502) were replaced by a *tetRA* cassette to construct the ΔP_*fliA*_::*tetRA* and ΔP_*flgM*_::*tetRA* mutants, respectively. The class 2 P_*flhB*_ promoter (region from 2023494 to 2022815) and class 3 P_*fliC*_ promoter (region from 2061043 to 2060527) were used as representative class 2 and class 3 flagellar promoters, respectively. Single-copy transcriptional fusion of P_*fliC*_ promoter to the fluorescent protein Venus was made by cloning into the plasmid pVenus using KpnI and EcoRI restriction sites and integrating the plasmids into the chromosome using the CRIM method. The P_*flhB*_ promoter (region from 2023494 to 2022815) was cloned into the plasmid pKW667 (35), containing the *mCherry* gene, using the XhoI and EcoRI restriction sites, yielding the plasmid pPROTet-*flhB*’-mCherry. The chloramphenicol resistance gene, P_*flhB*_ promoter, *mCherry* and terminator were then PCR amplified from the plasmid using the primers containing 40 base-pair homology to the flanking regions of the *araB* gene. The PCR product was used to replace *araB* gene with P_*flhB*_-*mCherry* reporter construct into the chromosome using λ-Red recombination (36). The integrated plasmids were then moved into the wild type and the different mutants by P22 transduction.

**Table 1:**
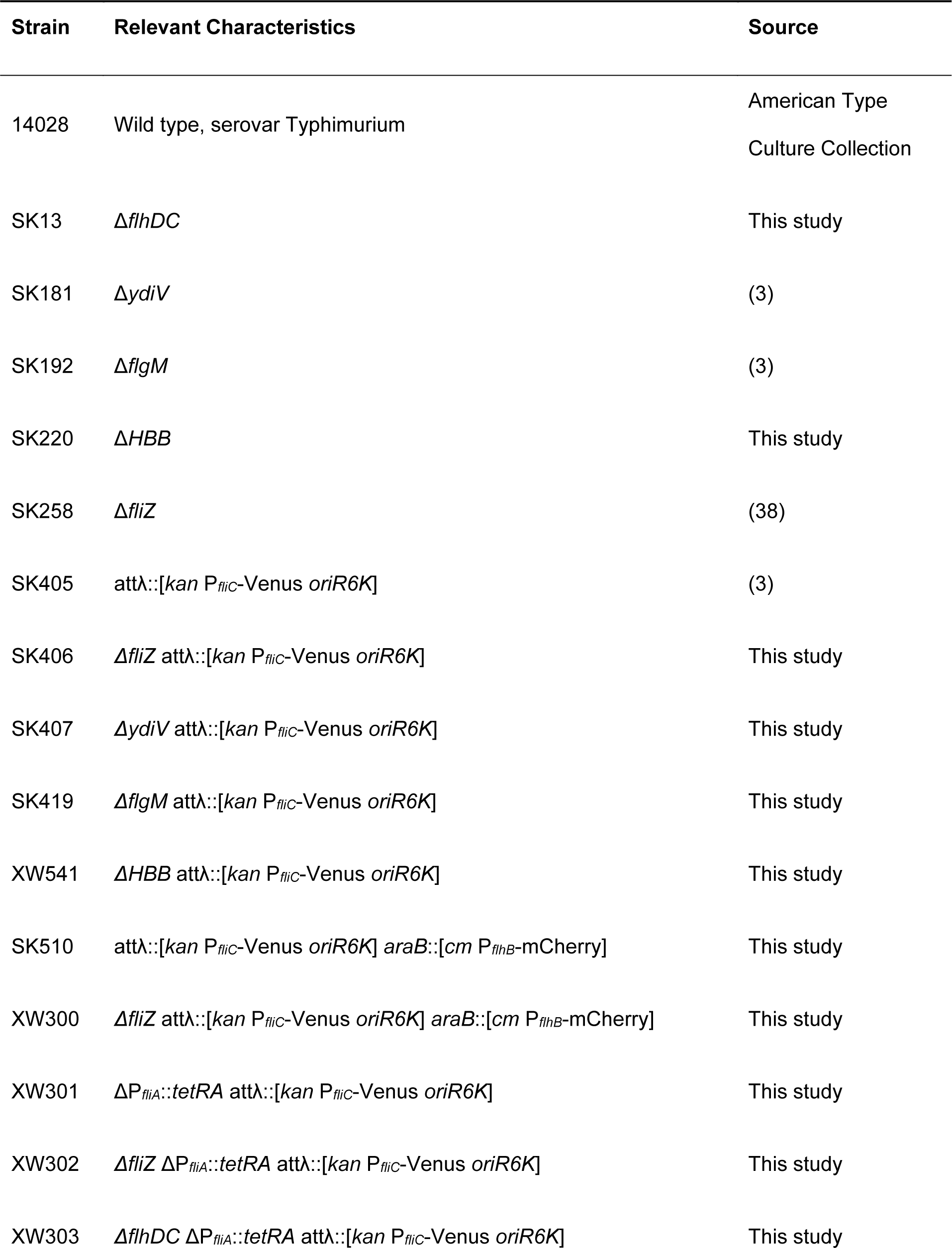
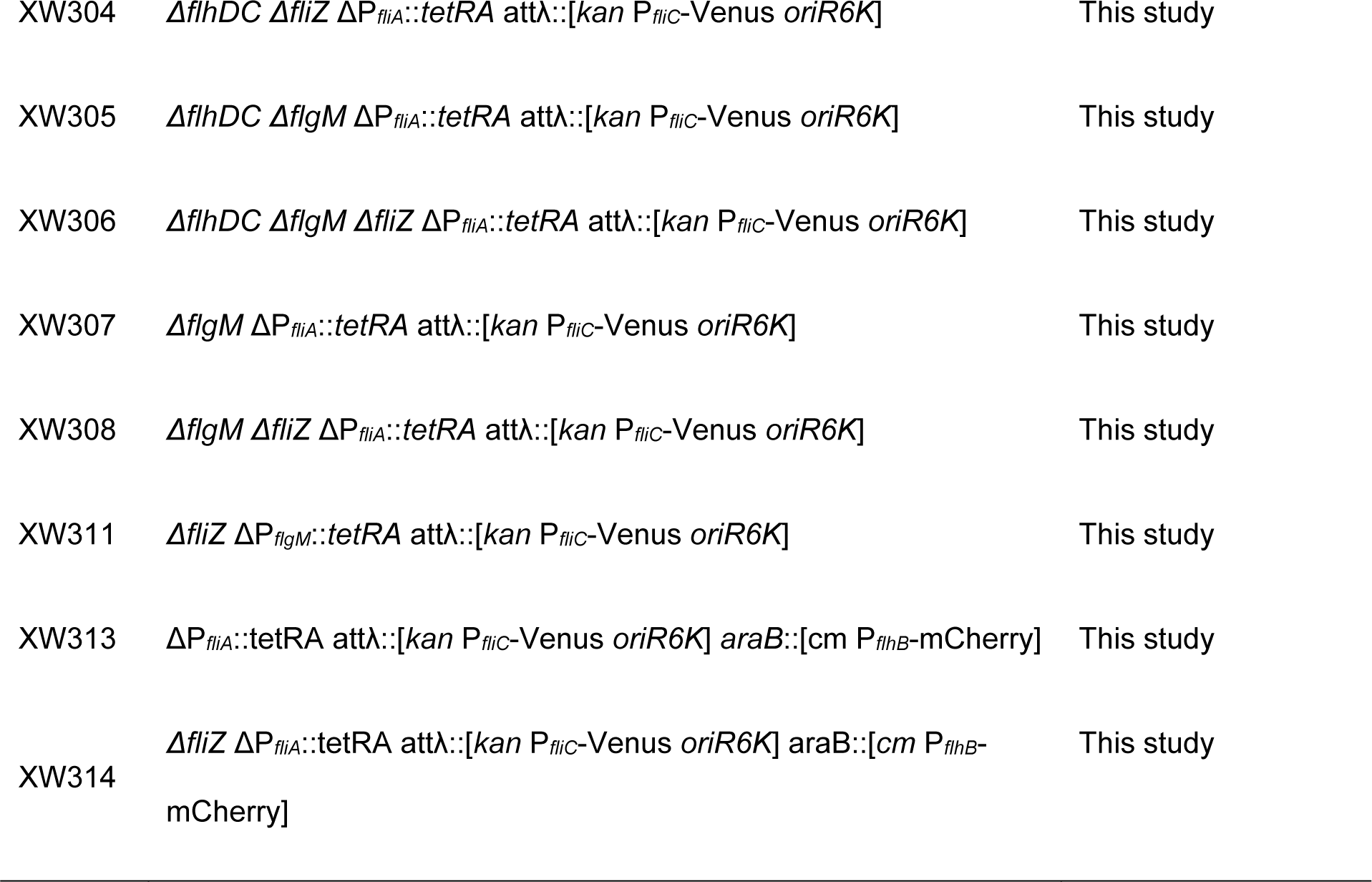
Strains used in this study

### Flow cytometry

Cells were grown overnight at 37°C in VBE medium supplemented with 0.2% glucose and 0.2% yeast extract as described previously (3, 37). Briefly, the cells were then subcultured to an optical density (OD_600_) of 0.05 in fresh VBE media supplemented with 0.2% glucose and the specified concentration of yeast extract and anhydrotetracycline (aTc). Following subculture, the cells were then allowed to grow at 37°C for 5 hours before harvesting. The cells were then pelleted by centrifuging at 3200 × *g* for 10 minutes and resuspended in phosphate-buffered saline (PBS) solution with 14.3 μM DAPI (4’-6-diamidino-2-phenylindole) and 50 μg/ml chloramphenicol. The suspension was then incubated at room temperature for half an hour. The cells were then analyzed using BD LSR Fortessa flow cytometer. Fluorescence values for approximately 100,000 cells were recorded using Pacific Blue channel (excitation: 405 nm; emission: 450/50 nm) for DAPI, fluorescein isothiocyanate channel (excitation: 488 nm; emission: 530/30 nm) for Venus fluorescence and phycoerythrin-Texas Red channel (excitation 561 nm; emission 610/20 nm) for mCherry fluorescence. The cells were distinguished from other debris by gating the population stained with DAPI. Data extraction and analysis for the FACS experiments were done using FCS Express Version 5 (De Novo Software). The data was exported to Microsoft Excel and further processed in Origin Pro 2018b to obtain histograms (for a single promoter) and density plots (for two promoters). The histograms show the distribution of promoter activities in individual cells as determined based on Venus fluorescence. The density plots show the distribution of promoter activities in individual cells as determined based on Venus and mCherry fluorescence. Data were smoothed and normalized to a peak value of 100 using the built-in function in FCS Express Version 5 to facilitate interpretation. All experiments were performed at least three times. Representative histograms and heatmaps are shown.

### Model of σ^28^/FlgM checkpoint

The model is a simplified version of a previously published model of the σ^28^/FlgM regulatory circuit (19). In particular, it does not include any other regulatory components besides σ^28^ and FlgM. Our rational here is to demonstrate that these two proteins are sufficient to generate class 3 bistablity. In addition, our analysis only focuses on the steady-state behavior of the flagellar network. While the model is formulated as a set of coupled differential equations, our subsequent analysis considered only the steady-state behavior as the corresponding experiments only measure gene expression at a single time point during exponential growth. In addition, we assumed that the associated between σ^28^ and FlgM is fast and effectively irreversible. Finally, we assumed that the degradation and dilution rates for species were the same: relaxing this assumption had no significant effect. Since we ignored temporal dynamics in our simulations, the associated kinetic parameters were taken to be one.

The governing equations for the model are:

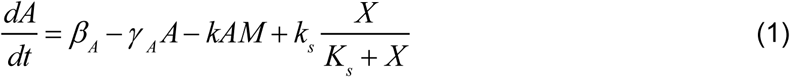

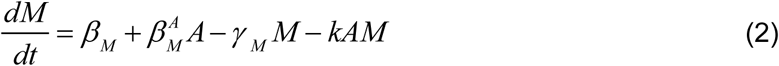

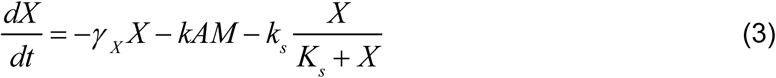

where *A* is the concentration of free σ^28^, *M* the concentration of free FlgM, and *X* is the concentration of σ^28^-FlgM complex.

The simulation results shown in Figure 4B plot the steady-state concentration of free σ^28^ (*A*) as a function of the secretion rate *k*_*s*_ for different values of basal FlgM expression *β*_*M*_. In simulation results shown in Figure 4C, we assumed that the secretion rate *k*_*s*_ was normally distributed in the population with varying means (〈*k*_*s*_〉 = 3,3.5,4,4.5,5) and fixed variance (var(*k*_*s*_=0.25). This was used to model the expected variability in the steady-state number of HBB’s within individual cells. In addition, we also added noise to calculated free σ^28^ concentrations to more accurately capture our single-cell gene expression experiments (a log-normally distributed random variable with zero mean and variance of 0.04 were added to the model results). The histograms result from Monte-Carlo simulations involving 5 million cells. In other words, we randomly sampled *k*_*s*_ from a lognormal distribution 5 million times with different mean values and then calcualted the associated free σ^28^ concentrations, with some additional noise added for aesthetic purposes (otherwise, the histogram is spiky at low σ^28^ concentrations). All simulations were performed in MATLAB (Mathworks, Natick, MA).

